# Genetically Determined Strength of Natural Killer Cells is Enhanced by Adaptive HLA class I Admixture in East Asians

**DOI:** 10.1101/2020.07.29.227579

**Authors:** Zhihui Deng, Jianxin Zhen, Genelle F. Harrison, Guobin Zhang, Rui Chen, Ge Sun, Qiong Yu, Neda Nemat-Gorgani, Lisbeth A. Guethlein, Liumei He, Mingzhong Tang, Xiaojiang Gao, Siqi Cai, Jonathan A. Shortt, Christopher R. Gignoux, Mary Carrington, Hongyan Zou, Peter Parham, Wenxu Hong, Paul J. Norman

## Abstract

Human natural killer (NK) cells are essential for controlling infection, cancer and fetal development. NK cell functions are modulated by interactions between polymorphic inhibitory killer cell immunoglobulin-like receptors (KIR) and polymorphic HLA-A, -B and -C ligands expressed on tissue cells. All *HLA-C* alleles encode a KIR ligand and contribute to reproduction and immunity. In contrast, only some *HLA-A* and *-B* alleles encode KIR ligands and they focus on immunity. By high-resolution analysis of *KIR* and *HLA-A*, *-B* and *-C* genes, we show that the Chinese Southern Han are significantly enriched for interactions between inhibitory KIR and HLA-A and -B. This enrichment has had substantial input through population admixture with neighboring populations, who contributed *HLA class I* haplotypes expressing the KIR ligands B*46:01 and B*58:01, which subsequently rose to high frequency by natural selection. Consequently, over 80% of Southern Han *HLA* haplotypes encode more than one KIR ligand. Complementing the high number of KIR ligands, the Chinese Southern Han *KIR* locus combines a high frequency of genes expressing potent inhibitory KIR, with a low frequency of those expressing activating KIR. The Southern Han centromeric *KIR* region encodes strong, conserved, inhibitory HLA-C specific receptors, and the telomeric region provides a high number and diversity of inhibitory HLA-A and -B specific receptors. In all these characteristics, the Southern Han represent other East Asians, whose NK cell repertoires are thus enhanced in quantity, diversity and effector strength, likely through natural selection for resistance to endemic viral infections.

## Introduction

Human leukocyte antigen (HLA) class I molecules are critical components of immunity, whose extreme variation associates with resistance and susceptibility to infection, multiple immune-mediated diseases and some cancers (Dendrou et al. 2018). *HLA class I* genes are located in the *major histocompatibility complex* (*MHC)* of chromosome 6 and encode proteins that bind peptide fragments derived from intracellular protein breakdown and transport them to the cell surface. In doing so they can communicate to the adaptive immune system’s T cells whether a tissue cell is healthy, or unhealthy due to infection or cancer. Subsets of HLA class I allotypes additionally contain an externally facing amino acid motif that binds killer cell immunoglobulin-like receptors (KIR), facilitating interaction with natural killer (NK) cells of innate immunity.

KIR are expressed on the surface of NK cells and regulate their functions through binding to HLA class I ligands on other cells (Cooper et al. 2009; Long et al. 2013). The functions of these interactions are crucial in immunity to aid recognition and elimination of infected or tumorous tissue, and in reproduction to regulate placentation and fetal development (Parham and Moffett 2013). In accordance with these critical and independent roles in human health, KIR and their HLA class I ligands are subject to natural selection, mediating their exceptional diversity across individuals, populations and species (Parham and Moffett 2013; Prugnolle et al. 2005). Indeed, *KIR* and *MHC* are some of the fastest evolving genomic loci in higher primates (Guethlein et al. 2015). Correlating with direct impact on both NK cell development and effector function (Freud et al. 2017; Vivier et al. 2011), numerous studies have implicated combinatorial diversity of *KIR* and *HLA class I* alleles with the course of specific infectious and autoimmune diseases, as well as the success of transplantation (Boudreau and Hsu 2018; Holzemer et al. 2017). Importantly, the quantity as well as quality of these interactions can influence individual responses to infection (Boelen et al. 2018; Pelak et al. 2011). Thus, the polymorphism of *KIR* and *HLA class I* has profound impact on human health. Under-explored are the scale and characteristics of *KIR* and *HLA class I* combinatorial diversity worldwide, and the processes that shape this diversity.

NK cells express overlapping subsets of KIR that are acquired stochastically during their development (Andersson et al. 2009). During this process, the interaction of inhibitory KIR with HLA class I KIR ligands broadens and strengthens subsequent effector functions of the NK cell repertoire (Bjorkstrom et al. 2016; Hoglund and Brodin 2010; Saunders et al. 2015). This education process matures some NK cells, allowing them to respond effectively to specific instances of infection or cancer, and enhances the NK cell repertoire compared to those that develop using other more conserved pairs of ligands and receptors. In this role, and also in pregnancy where HLA-A and -B have no function, HLA-C is dominant because all expressed HLA-C are KIR ligands (Guethlein et al. 2015). Four mutually exclusive sequence motifs define the four HLA class I epitopes that are KIR ligands: C1 is carried by subsets of HLA-C and HLA-B allotypes. C2 is carried by the other allotypes of HLA-C. Bw4 is carried by subsets of HLA-A and -B allotypes. The A3/11 motif is carried by a subset of HLA-A allotypes (HLA-A*03 and A*11). Thus, only some HLA-A and -B allotypes are KIR ligands and their main role is likely to diversify the NK cell response to pathogens.

The *KIR* locus on chromosome 19q13.4 varies in gene content, containing up to eight genes encoding inhibitory KIR and five encoding activating KIR (Wilson et al. 2000). Four of the inhibitory KIR and four activating KIR have well-characterized HLA-A, -B or -C ligands. Two broad groups of *KIR* haplotypes are present in every human population. *KIR A* haplotypes carry all four of the HLA-class I specific inhibitory receptors and are associated with resistance to infectious diseases (Bashirova et al. 2006). *KIR B* haplotypes are more variable in their gene number, carrying two or more genes for inhibitory receptors as well as various activating receptor genes, and favor fetal development (Parham and Moffett 2013). A recombination hotspot separates the *KIR* locus into two segments (Wilson et al. 2000). Two inhibitory receptors specific for HLA-C are encoded in the centromeric region, and two HLA-A and -B specific receptors are encoded in the telomeric region. Additional to gene content variation, polymorphism of both receptors and ligands can directly affect NK cell activity (Guethlein et al. 2015). Thus, by varying the number, density, specificity, strength or signaling properties of the receptor-ligand interaction, genetic variation of *KIR* and *HLA class I* can pre-determine functional differences in NK cell repertoires between individuals. This genetic diversity is substantial among populations, as demonstrated with high-resolution studies (Guethlein et al. 2015; Nemat-Gorgani et al. 2018). In such detailed analysis, Asian populations are under-represented.

Comprising 20% of the human population, the Chinese Han are the largest ethnic group in the world (Abdulla et al. 2009). The Han have a complex population history and are presently structured with the Northern and Southern Han forming two main subgroups that are separated geographically by the Yangtze River (Wen et al. 2004). The Southern Han originated through large scale population movement from the north ~1500 years ago, in parallel with admixture with resident and neighboring populations (Hellenthal et al. 2014; Wen et al. 2004). Importantly for the current study, the major genetic distinction between the Northern and Southern Han occurs in the *MHC*, and localizes to the region that spans *HLA-A*, *-B* and *-C* (Chen et al. 2016; Xu et al. 2009). The most significant component of this difference is the *A*33:03-B*58:01-C*03:02 HLA class I* haplotype, which is common in the Southern Han and remains conserved across multiple unrelated individuals (Chen et al. 2016). Such strong linkage disequilibrium is consistent with recent acquisition of this haplotype by admixture (Chen et al. 2016). This haplotype encodes two KIR ligands, HLA-B*58:01 and C*03:02 (Guethlein et al. 2015). Although less is known of *KIR* allele diversity in the Han, several studies established that the genes characteristic of *KIR A* haplotypes are common, and demonstrated differences in their distribution among the different Han groups and among other resident populations (Bao et al. 2013; Wang et al. 2012; Yao et al. 2011). These studies also confirmed that *KIR* and *HLA class I* combinatorial diversity is an important factor in pregnancy syndromes, infectious disease, blood cancers and transplantation outcome in the Han. They also uncovered both similarities and differences from the specific disease associations observed in Europeans (Bao et al. 2016; Jiang et al. 2013; Long et al. 2015; Shen et al. 2016; Su et al. 2018). To investigate these findings, we have examined how demographic and evolutionary processes have shaped combinatorial diversity of HLA class I and KIR in the Chinese Southern Han.

## Materials and methods

### Study samples

Peripheral blood samples were collected from 306 unrelated healthy volunteer blood donors from Shenzhen, Guangdong, China. All donors self-identified to be of Han ethnicity from southern China. All subjects provided written informed consent for participation in the present research, which was approved by the ethics review board of Shenzhen Blood Center, Shenzhen, Guangdong, China.

### Genomic DNA extraction

Genomic DNA was extracted from 400 μl of peripheral blood using a MegCore Nucleic Acid Extractor (MegCore, Taiwan, China). DNA purity and concentration were tested by UV-spectrophotometry using a Biophotometer (Eppendorf, Hamburg, Germany) and adjusted to a concentration of 50-100 ng/μl.

### High-resolution HLA-A, -B and -C genotyping

*HLA-A*, *-B* and *-C* genotyping was performed using the AlleleSEQR HLA sequencing-based genotyping commercial kit (Atria Genetics, San Francisco, USA). According to the manufacturer’s instructions, exons 2-4 for *HLA-A*, *-B* and *-C* were sequenced in both directions using an ABI 3730XL DNA sequencer (Applied Biosystems, Foster City, CA, USA). HLA genotypes were assigned using the Assign 4.7 software (Conexio Genomics, Fremantle, Australia). Samples giving ambiguous allele combinations by sequencing were further resolved using HLA PCR-SSP (Olerup, Stockholm, Sweden).

### High-resolution KIR genotyping

The presence or absence of *KIR2DL1*, *2DL2*/*3*, *2DL4*, *2DL5*, *2DS1*, *2DS2*, *2DS3*, *2DS4*, *2DS5*, *3DL1*/*S1*, *3DL2* and *3DL3* was first determined for each individual using the ‘KIR Ready Gene’ PCR-SSP kit (Inno-Train Diagnostik GmbH, Frankfurt, Germany). The *KIR* genes identified using PCR-SSP were then subject to nucleotide sequencing of all exons (Deng et al. 2018). Sequencing reactions were performed using ABI PRISM BigDye Terminator Cycle Sequencing Ready reagents and analyzed using an ABI 3730 DNA Sequencer (Applied Biosystems, Foster City, USA). *KIR* alleles were assigned using Assign 4.7 allele identification software (Conexio Genomics, Fremantle, Australia), and release 2.6.1 (February 2015) of the Immuno-Polymorphism database (IPD) (Robinson et al. 2015). When the sequencing results gave ambiguous allele combinations, we used group-specific PCR primer pairs to amplify and sequence the target alleles separately (Zhang and Deng 2016)

### Novel KIR alleles

To confirm and fully characterize any novel allele identified during amplicon sequencing we cloned and sequenced *KIR* transcripts. Further samples of peripheral blood samples were collected, and total RNA isolated using the Maxwell 16 low elution volume simplyRNA Blood Kit (Promega, Madison, USA). Complementary DNA (cDNA) was synthesized using the Transcriptor First Strand cDNA Synthesis Kit (Roche, Basel, Switzerland). *KIR* transcripts were amplified specifically from cDNA using primer pairs described previously (Yawata et al. 2006), with addition of KIR3DL3-specific primers (forward 5’-GGTTCTTCTTGCTGGAGGGGC-3’ and reverse 5’-TTACACGCTGGTATCTGTTGGGG-3’). The amplified transcripts were cloned using the TA cloning kit (Takara, Dalian, China) and at least three clones of any novel allele were sequenced. The sequences of novel *KIR* alleles were submitted to GenBank and the IPD KIR database (Robinson et al. 2015) to obtain official names.

### Admixture Estimates

Whole genome SNP genotypes for Japanese (N = 104), Vietnamese (N = 99), Han from Beijing (N = 103), Southern Han (N = 105), and Dai (N = 93) were obtained from the 1000 Genomes Project (Auton et al. 2015). We used any SNPs having minor allele frequency >1% and independent of other SNPs (linkage disequilibrium, r^2^ < 0.3). Admixture was calculated for chromosome 6 using the ADMIXTURE program (Alexander et al. 2009), with the unsupervised option and k=3. Two regions were analyzed, the *MHC* (chr6:28,477,797–33,448,354: 3,541 SNPs) and chromosome 6 excluding the *MHC* (84,898 SNPs). We selected a K of 3 to represent the three primary ancestry groups in the region that are represented in the 1000 Genomes data: Japanese, South East Asian, and East Asian (Chen et al. 2016). *HLA class I* alleles were obtained from the 1000 Genomes Project data (Gourraud et al. 2014). We analyzed the Hondo Japanese (JPT), Vietnamese (KHV), Chinese Dai (CDX), Chinese Southern Han (CHS), and Beijing Han (CHB) Validating their use for this purpose, the correlation of the *HLA class I* allele frequencies between our study population and the CHS is 0.95 (p= 6.65^−11^, Figure S1). Individuals were considered carriers if they had at least one copy of the respective allele. Distributions of ancestry proportions for carriers and non-carriers of specific *HLA* alleles were compared using a Wilcoxon test, using the wilcox.test function in R (R Development Core Team 2008).

### Estimates of nucleotide diversity

We used π (Nei and Takahata 1993) to measure the nucleotide diversity of haplotypes carrying specific *HLA-B* alleles. We used the phased genomes of the Chinese Southern Han (CHS) population available from the 1000 Genomes Project (Auton et al. 2015), and extracted the genomic region containing the *HLA-B* and *-C* genes, with 500kbp flanking on each side. For each carrier of a given allele, we identified (by sequence) and retained the haplotype representing the allele of interest. For each given allele, we pooled all of the respective haplotypes present in the population and calculated π in 100bp windows using VCFtools (Danecek et al. 2011). Distributions of π values were compared between respective alleles with a Wilcoxon test using the wilcox.test function in R.

### Tests for positive selection affecting specific HLA class I alleles

We filtered 1,000 Genomes genotyping data of chromosome 6 from the CHS population to remove non-biallelic and duplicated SNPs (Purcell et al. 2007), then phased using the program Eagle (Loh et al. 2016). We used the program Selscan (Voight et al. 2006) to calculate integrated haplotype statistic (iHS). The statistic is a measure of haplotype diversity associated with a given genetic variant, where lower diversity and longer haplotypes correlate with selection of that variant.

To determine if specific *HLA class I* alleles have been targeted by directional selection in the Chinese Southern Han we again used the 1,000 Genomes SNP data from the CHS population. SNPs within the following hg19 coordinates were used: *HLA-A*, Chr6: 29,910,089 – 29,913,770; *HLA-B*, Chr6: 31,321,648 – 31,325,007; *HLA-C,* Chr6: 31,236,517 – 31,239,917. We phased haplotypes from individuals positive for each given *HLA class I* allele and aligned them to reference sequences to identify the haplotype containing that allele. The alignments were then used to identify ‘tagging’ SNPs that could be used to identify each given *HLA class I* allele. The criteria for choosing tagging SNP alleles were that they must be present in every individual carrying the corresponding *HLA class I* allele and that they must be absent from the other *HLA class I* alleles in the analysis. We analyzed the alleles present on the 10 most frequent *HLA class I* haplotypes that we observed in the Chinese Southern Han; *HLA-B*15:02* was excluded because we were not able to identify unique tagging alleles on haplotypes carrying this allele. For each tagging SNP, we calculated the integrative haplotype score (iHS) using SelScan. We used the absolute value of iHS since derived alleles under selection will have a negative value and ancestral alleles under selection will have a positive value (Szpiech and Hernandez 2014). Using a Wilcoxon two-sample test, we examined whether the distributions of absolute iHS values differed between tagging SNPs of *HLA* alleles and SNPs of the full chromosome 6.

### Haplotype and ligand frequencies

*KIR* and *HLA-A*, *-B* and *-C* allele frequencies were calculated from the observed genotypes. For individuals genotyped as homozygous for an allele of a given *KIR* that has presence/absence polymorphism, the number of copies present was determined by analyzing LD with alleles of the flanking genes. The subsequent genotype distributions for all loci were consistent with Hardy-Weinberg equilibrium. *KIR* haplotype frequencies were determined using PHASE II (Stephens and Donnelly 2003). The following parameters were used; -f1, -x5, and -d1, and from the output, the two haplotypes with highest probability were taken for each individual. Watterson’s homozygosity F test was performed using Pypop software (Lancaster et al. 2007), with 10,000 replicates to calculate the normalized deviate F_nd_ test (Salamon et al. 1999).

*HLA class I (A-B-C)* haplotype frequencies were determined using the EM algorithm of ‘Arlequin software version 3.5 (Excoffier and Lischer 2010). For comparison with other populations, we only used populations for which *HLA class I* genotype data were available from every individual sampled, and for which the resolution of genotyping was the same as the Southern Han described here. We therefore used the subset of populations described in the 13^th^ International Histocompatibility Workshop and Conference report that have 50 or more *HLA-A*, *-B* and *-C* genotyped individuals (Meyer et al. 2007). These were supplemented with our own data from the Ga-Adangbe from Ghana in West Africa (Norman et al. 2013), KhoeSan from Southern Africa (Nemat-Gorgani et al. 2018), Yucpa from South America (Gendzekhadze et al. 2009), Europeans from the USA (Norman et al. 2016) and Hondo Japanese (Yawata et al. 2006). We compared the proportion of *HLA class I* haplotypes encoding one KIR ligand to those encoding more than one KIR ligand across populations using a two-proportions Z test, using the prop.test function in R (R Development Core Team 2008).

### Comparison of HLA class I and KIR ligand distributions

Clustering based on allele frequencies: Any *HLA class I* allele occurring in fewer than two of the populations studied was excluded from this analysis. The allele frequencies of all three *HLA class I* genes were used for each population. Cluster dendrograms were constructed using R 3.4.3, hclust with 1,000 bootstrap values. The package used was fpc (Hennig 2020). Cluster dendrograms were constructed in the same manner using the frequencies of the *HLA class I* haplotypes encoding one, two or three KIR ligands.

### Assessment of receptor/ligand quality and quantity

As described previously (Nemat-Gorgani et al. 2018), experimental data were used to determine the interacting pairs of KIR and HLA class I, which are listed in **Figure S2**. To determine the quantity of receptor/ligand interactions, the number of KIR/HLA allotype pairs that are known to interact were summed for each individual, and homozygous KIR or HLA allotypes were counted twice. To determine the diversity of interactions, the number of different KIR/HLA allotype pairs that are known to interact were summed for each individual (in this case homozygous allotypes were counted once). Populations were compared using unpaired t tests, using GraphPad software.

## Results

### High Frequency of KIR ligands in the Southern Han

All HLA-C and subtypes of HLA-A and -B allotypes are ligands for KIR, which are expressed on the surface of NK cells to modulate their functions in immunity and reproduction. Within human populations, *HLA class I* haplotypes tend to form a balance between those that encode HLA-A or -B KIR ligands and those that do not (Guethlein et al. 2015). To determine if this pattern is also observed in the Chinese Southern Han, we analyzed the *HLA-class I* genes of 306 healthy individuals. We identified 27 *HLA-A*, 54 *HLA-B* and 29 *HLA-C* alleles (Figure S3). Each of these 110 alleles encodes a different HLA class I allotype, and 58 of them are known KIR ligands (Figure 1A). The majority of 233 *HLA class I* haplotypes, including the ten most frequent (Figure 1B), encode more than one KIR ligand (70.3% of distinct haplotypes; 81.8% by frequency, Figure S4A). This observation is unusual and indicates the balance between having and not having KIR ligands at HLA-A and -B is perturbed in the Chinese Southern Han.

**Figure 1.**
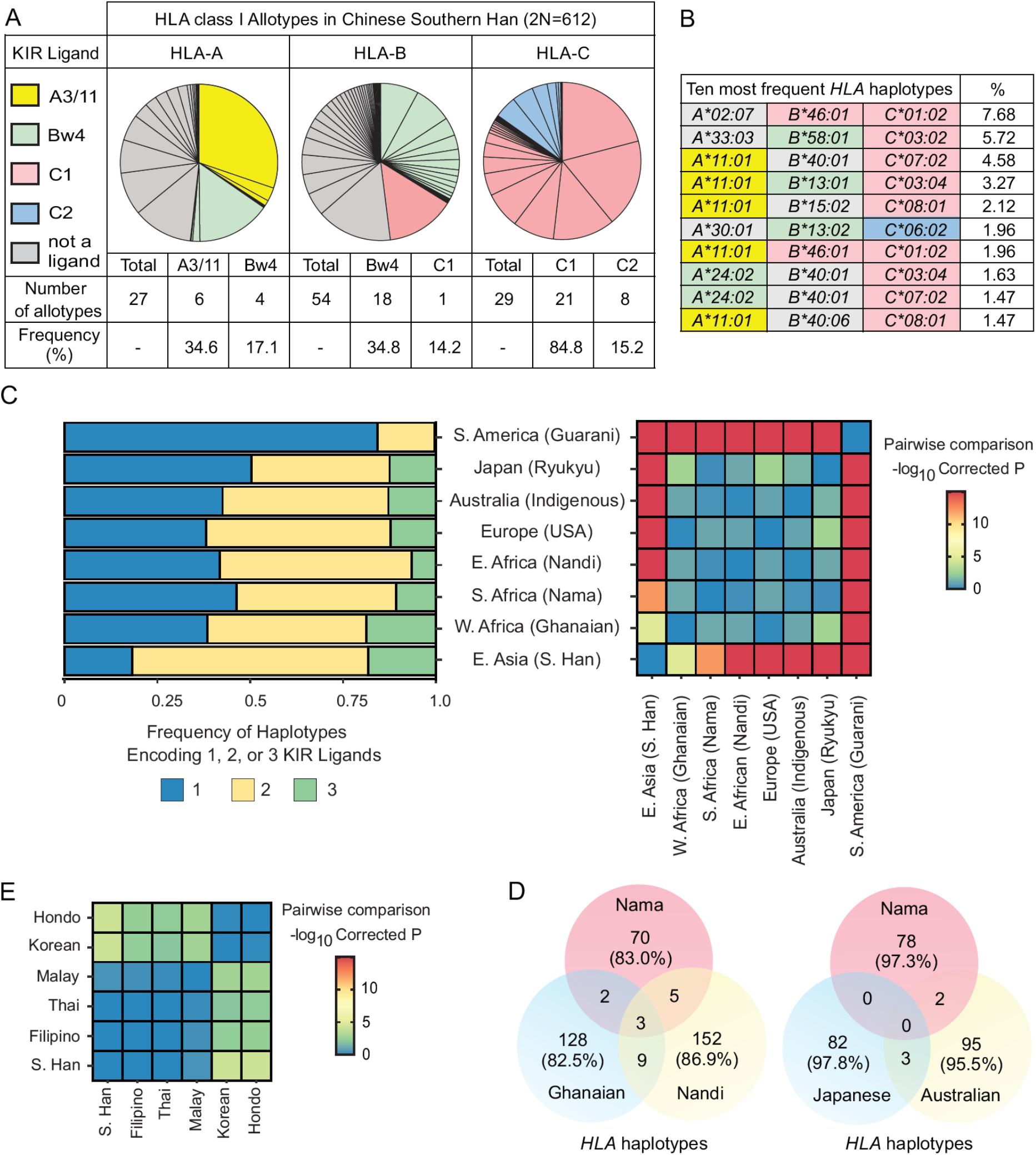
Chinese Southern Han *HLA class I* haplotypes express multiple KIR ligands. A. Pie charts show the frequency spectra for HLA-A, -B and -C allotypes of the Southern Han cohort of 306 unrelated individuals (2N=612). Each pie segment represents one allotype. Alternative sequence motifs in the α1 domain of the HLA class I molecule determine the four epitopes recognized by different KIR, and which are also called KIR ligands. The A3/11 epitope is carried by HLA-A3 and -A11 (yellow colored pie segments); the Bw4 epitope is carried by subsets of HLA-A and -B allotypes (green colored pie segments). The C1 epitope is carried by a majority of HLA-C allotypes, as well as by HLA-B*46 and HLA-B*73 (red colored pie segments). The C2 epitope is carried by all HLA-C allotypes that do not carry C1 (blue-colored pie segments). Grey-colored pie segments correspond to allotypes that are not KIR ligands. Figure S2 lists all the HLA-A, -B and -C allotypes present in the study population and shows which KIR ligand motifs they carry. B. Shows the ten most frequent *HLA class I* haplotypes in the Southern Han and their frequencies (2N=612). Colored shading indicates *HLA class I* alleles that encode KIR ligands, as described in panel A. C. (left) Bars show the combined frequencies of *HLA class I* haplotypes encoding one (blue), two (gold) or three (green) KIR ligands in eight representative populations worldwide (Southern, Western and Eastern Africa, Europe, Oceania, South America, Japan and Chinese Southern Han). (Right) Heat-plot shows pairwise comparisons between populations of the proportion of *HLA class I* haplotypes encoding one KIR ligand to those carrying two or more KIR ligands. Colors correspond to −log^10^ of a Benjamini-Hochberg corrected p, as shown in the key. D. Heat-plot shows pairwise comparisons of Chinese Southern Han with five East/ South East Asian populations of the proportion of *HLA* haplotypes encoding one KIR ligand to those carrying two or more KIR ligands. Colors correspond to −log_10_ of a Benjamini-Hochberg corrected p, as shown in the key. E. Venn diagrams show the distribution of *HLA class I* haplotypes within representative subsets of populations. The number of haplotypes in each overlapping region is given. The % values indicate the combined frequency of haplotypes unique to a population when compared to the other populations in the diagram.

To investigate the unusually high frequency of KIR ligands, we compared Southern Han *HLA class I* haplotypes with those of sub-Saharan African, Oceanian, European and South American populations that represent major modern human groups (Rosenberg et al. 2002; Tishkoff et al. 2009). In this data set, rather than the larger and more widely studied Hondo Japanese population, a Ryukyu Japanese population was included because they more closely represent the Japanese population prior to admixture with Han (Takeuchi et al. 2017). Among the eight populations, 1,034 different *HLA class I* haplotypes were observed (Figure S4B). Six populations have a similar distribution of KIR ligands, with each population having an approximately equal frequency of *HLA class I* haplotypes carrying one and two KIR ligands, and a smaller frequency of haplotypes carrying three KIR ligands (Figure 1C). Only Southern Han and South Americans differed from this pattern, with the Han encoding more and the Amerindians encoding less KIR ligands per haplotype than other populations (Figure 1C). The difference in the proportion of *HLA class I* haplotypes encoding one versus more than one KIR ligand between the Southern Han and each of the other seven representative populations is statistically significant, as is that between Amerindians and the other populations (Two-proportions Z-test, Benjamini-Hocherg corrected p <0.001, Figure 1C). The allele frequency distribution of South American Amerindians was likely influenced by severe population bottlenecks, leading to a reduced genome-wide diversity compared with other populations (Fagundes et al. 2008; Raghavan et al. 2015), whereas the Han were not subject to severe population-specific bottleneck (Henn et al. 2012; Lu et al. 2016; Schiffels and Durbin 2014). To examine if the Chinese Southern Han are representative of other related populations, we examined groups from East Asia (Hondo Japanese and Korean) and Southeast Asia (Thai, Malay and Filipino). This analysis showed these populations also have a high frequency of *HLA class I* haplotypes encoding multiple KIR ligands (Figure 1D). Our analysis thus shows that East Asian and South East Asian *HLA class I* haplotypes encode more ligands for inhibitory KIR than the haplotypes of any other populations.

Despite having distinct population histories, the sub-Saharan African, Oceanic, European and Ryukyu Japanese populations all have a similar mean number of KIR ligands per *HLA* haplotype (Figure 1C). However, very few *HLA class I* haplotypes are shared by any of these populations. For example, only 19 of 369 haplotypes detected in Africans are present in more than one of the three African populations studied, and comparing the disparate Southern African Nama, Indigenous Australian and Ryukyu Japanese populations revealed just five haplotypes in common (Figure 1E).

### The Chinese Southern Han acquired HLA haplotypes encoding multiple KIR ligands by admixture

Previous analyses suggested that specific *MHC* region haplotypes (which include the *HLA* genes) present in the Chinese Southern Han were obtained from the Northern Han through admixture (Chen et al. 2016). The most frequent HLA class I allotypes contributing to the enrichment of KIR ligands in the Chinese Southern Han are HLA-A*11, -A*24, -B*46, and -B*58 (Figure 1B). We therefore examined the relative contributions of admixture to the high frequency of these alleles in the Chinese Southern Han. For this analysis we considered known admixture events (Hellenthal et al. 2014; Wen et al. 2004; Xu et al. 2009) and drew upon the 1000 Genomes SNP and *HLA* genotype data (Auton et al. 2015; Gourraud et al. 2014) from Hondo Japanese, Vietnamese, Dai, Beijing Han, and Chinese Southern Han. Consistent with previous work examining whole-genome data (Takeuchi et al. 2017), in analyzing chromosome 6 we identified three primary genetic ancestries, corresponding to the Japanese, East Asian (Southern and Beijing Han) and South East Asian (Vietnamese and Dai) population groups (Figure 2A). That we identify a higher ‘Japanese’ component in the Beijing than Southern Han (36% vs 22%: Figure 2A) likely reflects the higher proportion in Beijing of Northern Han (Auton et al. 2015), a population from which we have no data for the current study. Supporting this observation, the greatest contribution from China to Japanese ancestry is from the Northern Han (Chen et al. 2016; Takeuchi et al. 2017).

**Figure 2.**
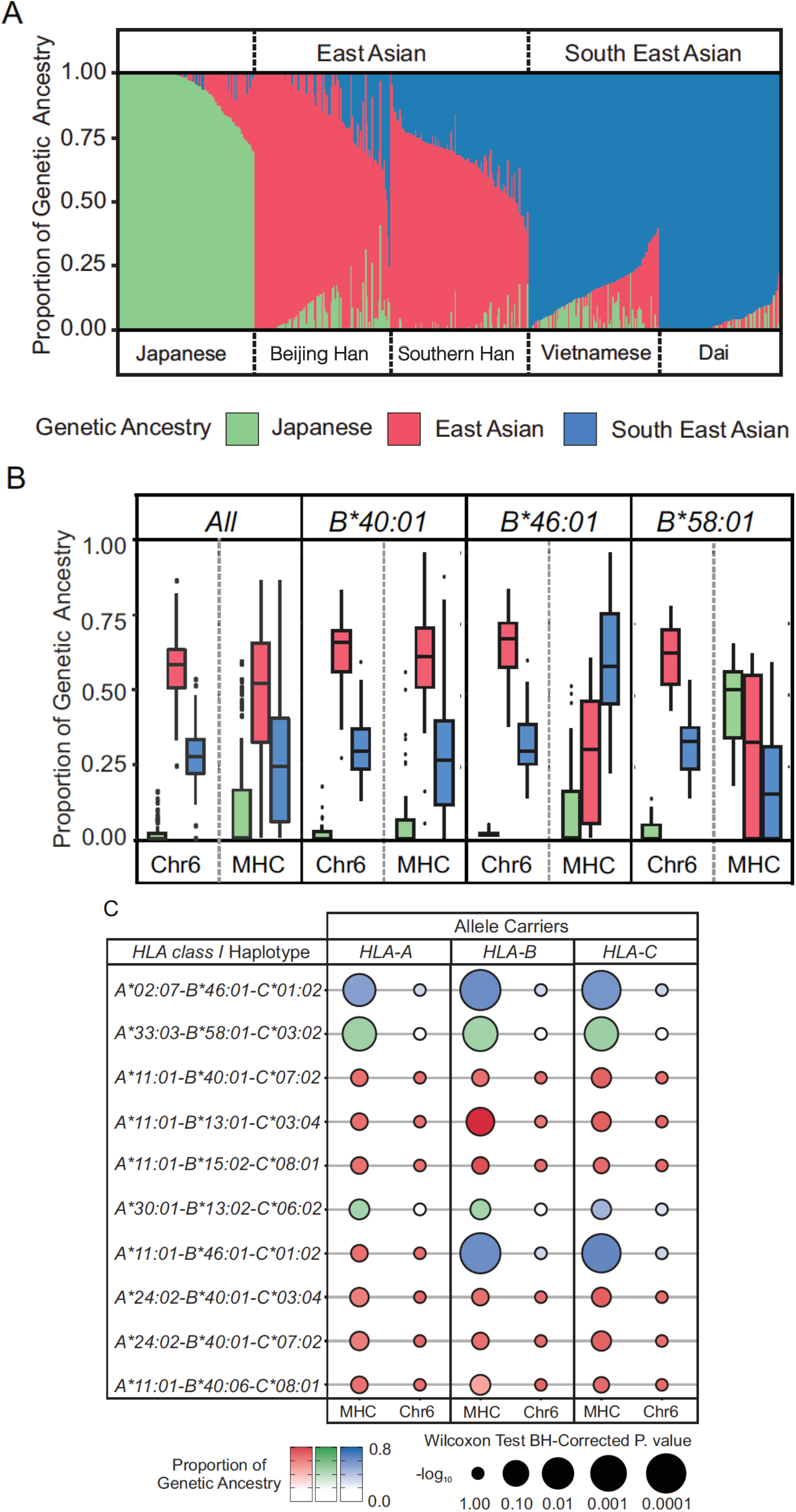
HLA-B*46:01 and -B*58:01 were acquired by admixture into the Chinese Southern Han. A. Shown are the relative proportions of genetic ancestry among Asian populations from the 1000 genomes project, plotted by considering three ancestral groups (K = 3: Japanese (green), East Asian (red) and South East Asian (blue)). B. Shown are the relative proportions for each of the three genetic ancestries in chromosome 6 (left) and within the *MHC* (right) for selected Chinese Southern Han individuals, shown from left to right; all individuals, *B*40:01* carriers, *B*46:01* carriers, *B*58:01* carriers. C. Shown for each of the ten most frequent *HLA class I* haplotypes in the Chinese Southern Han is a comparison of mean admixture proportion of the genetic ancestry group that is most abundant in *MHC* compared to the proportion of that ancestry in chromosome 6 with *MHC* excluded. The size of the circle represents the −log_10_ (Benjamini-Hochberg corrected p) from a Wilcoxon test. The difference in size and shade between the two circles corresponds to extent and direction, respectively, of any shift in the genetic ancestry proportions between the *MHC* and the remainder of the chromosome 6.

We compared the relative proportions of the three genetic ancestries in the Chinese Southern Han within the *MHC* region of chromosome 6 to their proportions throughout chromosome 6 excluding the *MHC*. This analysis revealed a predominance of East Asian ancestry throughout the length of chromosome 6, including the *MHC* (Figure 2B). In carriers of *HLA-B*40:01*, the most frequent *HLA-B* allele in Chinese Southern Han, there is also clear East Asian ancestry throughout the length of chromosome 6 (Figure 2B). The proportion of East Asian ancestry is similar in *B*40:01* carriers than non-carriers (Wilcoxon test, p=0.98). By contrast, among *HLA-B*46:01* carriers, the *MHC* is primarily of South East Asian ancestry (Figure 2B) with carriers having a significantly higher proportion of South East Asian genetic ancestry in the *MHC* than outside the *MHC* (p = 2.7^−6^), or within the *MHC* of non-carriers (p = 1.9^−5^). Similarly, among *HLA-B*58:01* carriers, the *MHC* region is primarily of Japanese ancestry (Figure 2B), with carriers having a significantly higher proportion of Japanese genetic ancestry within the *MHC* than outside the *MHC* (p = 2.4^−4^) and compared with non- *B*58:01* carriers (p = 6.1^−7^). Excluding the *MHC,* carriers of any of these three alleles show East Asian ancestry along chromosome 6 (Figure 2B). Further supporting the observed population structure as specific to the *MHC* region, among the three ancestral groups the F_ST_ values range from 0.098 – 0.161, compared with 0.012 – 0.017 for the remainder of chromosome 6.

Based on the analysis of *B*46* and *B*58*, we examined the proportions of genetic ancestry of alleles that comprise the 10 most frequent *HLA class I* haplotypes observed in the Chinese Southern Han. The primary genetic ancestry outside of the *MHC* region was determined as East Asian for every allele studied (Figure 2B). Thus, for our comparisons, we determined the primary genetic ancestry in the flanking *MHC* region for each allele and then determined the relative proportion of that ancestry in the remainder of the *MHC*. This analysis identified six haplotypes maintaining strong evidence of East Asian genetic ancestry both within the *MHC* and throughout chromosome 6. These haplotypes include those that carry *A*11:01* and *A*24:02*, as well as *B*40:01* (Figure 2C). It was shown previously that HLA-A*11 and -A*24 derive from introgression with archaic humans (Abi-Rached et al. 2011) and our results and others (Gonzalez-Galarza et al. 2015; Solberg et al. 2008) thus suggest these haplotypes are now endemic to East Asia. The analysis also identified four haplotypes having genetic ancestry within the *MHC* that is distinct from the ancestry of the remainder of the chromosome (Figure 2C). For three of the haplotypes, which include the two most frequent haplotypes in the population, this distinction is statistically significant (p_corr_ < 0.01). Two of these haplotypes contain *B*46:01* and one contains *B*58:01* (Figure 2C). In total, four of five of the *HLA-B* alleles that encode a KIR ligand and are present on these 10 most frequent haplotypes show increased evidence for admixture in the *MHC* region. By contrast, neither of the two *HLA-A* alleles that encode a KIR ligand show a genetic ancestry within the *MHC* that differed from the East Asian ancestry throughout chromosome 6. This finding suggests that the number of *HLA-B* genes encoding KIR ligands was enhanced in Chinese Southern Han by admixture with neighboring or displaced populations. In summary, these findings clearly show that the *B*46:01* and *B*58:01* alleles are present in Chinese Southern Han through admixture.

### Positive selection favors HLA haplotypes expressing more than one KIR ligand in Chinese Southern Han

To investigate whether or not the admixed haplotypes were also subject to natural selection we examined further characteristics of their diversity and distribution. We first measured nucleotide diversity (π) of the genomic sequence flanking +/− 500kb of specific *HLA-B* alleles (Figure 3A). We found significantly reduced nucleotide diversity of haplotypes containing *HLA-B*46* compared to haplotypes containing *HLA-B*40* (mean π of *B*40* = 2.2 × 10^−3^, *B*46* = 0.6 × 10^−3^, Wilcoxon test, p = 1.24 × 10^−12^). We also observed that haplotypes containing *B*58* have lower diversity than *B*40*, but this reduction was not statistically significant (mean π of *B*58* = 1.6 × 10^−3^, Wilcoxon test, p =0.12). This reduced diversity suggests that *B*46* haplotypes have arisen in frequency in the Chinese Southern Han without accumulating mutations. To further explore this finding, we used the iHS statistic, which identifies genomic variants that have increased in frequency recently and rapidly under natural selection, so that their haplotypic background has not yet been diversified by recombination (Voight et al. 2006). We identified a strong signal of recent selection (iHS >= 99^th^ percentile) that falls precisely in the *MHC* of the Chinese Southern Han (Figure 3B).

**Figure 3.**
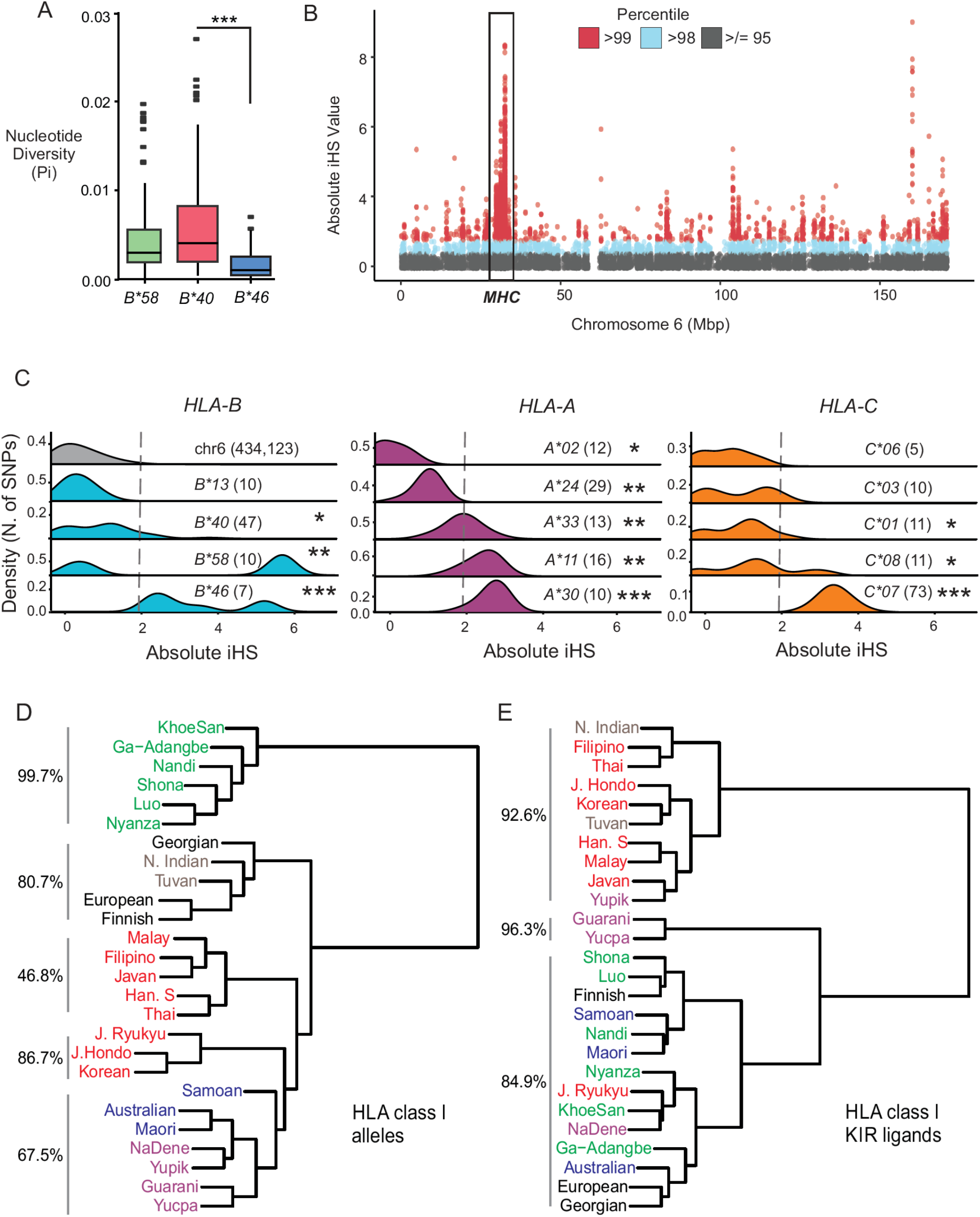
Positive selection has targeted *HLA class I* genes in the Chinese Southern Han. A. Shown is the nucleotide diversity (π) of genomic sequence +/− 500kbp of the *HLA-B* and *-C* genes for haplotypes containing the specific *HLA-B* alleles, *B*40*, *B*46* and *B*58*. π was measured in windows of 100bp. *** p <0.001 by Wilcoxon test. B. Manhattan plot shows the absolute iHS values above the 95^th^ percentile calculated for independent SNPs throughout chromosome 6 in Chinese Southern Han. The *MHC* region is boxed. C. Density plots show the distribution of absolute iHS values for chromosome 6 (top left, grey shading), and for SNPs unique to haplotypes carrying specific *HLA-B* (left, cyan), *HLA-A* (center, purple) and *HLA-C* (right, orange) alleles, as indicated in each plot. The number of SNPs unique to each of the *HLA* alleles is shown in brackets next to the allele name. For each allele, the distribution of iHS values was compared to the distribution on chromosome 6 using a Wilcoxon two-sample test (* p < 0.05, ** < 0.05^−5^, *** < 0.05^−10^). Grey dashed line marks the 95^th^ percentile of iHS values for chromosome 6 SNPs (=1.93). The density shown is the kernel density estimate of the SNP counts associated with the distribution of absolute iHS values. D. Shows cluster analysis of *HLA class I* allele frequencies from 26 populations. Vertical lines at the left show the clusters identified when five groups were specified in the input parameters (k=5) and the support (%) from 1,000 bootstrap replicates. Population names in green text indicate sub-Saharan African populations, red text – East Asian, brown text – Northeast Asian (Tuva) and South Asian (Indian), blue text – Oceanic, purple text – Amerindian (Yupik are North Amerindian who back-migrated to Siberia (Raghavan et al. 2015), NaDene are North American). J. Hondo are Japanese from the major islands of Japan, J. Ryukyu are Japanese from Okinawa. The *HLA class I* haplotypes detected in each population are described in Figure S4D. E. Shows cluster analysis of the combined frequencies of *HLA class I* haplotypes carrying 1, 2 or 3 KIR ligands. Vertical lines at the left show the clusters identified when three groups were specified (k=3) and the support (%) from 1,000 bootstrap replicates.

To investigate the patterns of selection specific to *HLA-B*46:01* and *B*58:01* haplotypes we first identified SNPs that characterize those haplotypes and then compared their distribution of iHS values to the distribution of all the SNPs of chromosome 6 (Figure 3C). For *B*46:01* the mean absolute iHS of 3.38, was significantly higher than the mean for chromosome 6 of 0.785 (Wilcoxon two-sample test, p= 5.77^−6^), as was the mean iHS for *B*58:01* (3.43, p= 1.8^−3^). Although the signal for *B*58:01* is weaker, there is a more distinct subset of SNPs having extremely high iHS values (>99^th^ percentile, Figure 3C), which could indicate recent selection of an older haplotype, although it was not possible from our analysis to determine if the SNP allele was ancestral or derived in each case. Interestingly, the mean iHS for *B*40:01* associated SNPs was also significantly higher than the chromosome average (1.88, p= 7.7^−6^). However, fewer *B*40:01* specific SNPs had an iHS value in the 95th percentile than *B*58:01* or *B*46:01* specific SNPs (30%, 50%, and 100% of SNPs respectively). Because the most frequent *B*40:01* containing haplotypes in the Chinese Southern Han carry either *A*11:01 or A*24:02,* which are KIR ligands (Figure 1B), we extended the analysis to these alleles (Figure 3C). Again, this analysis showed both *A*11:01* (mean = 2.8, Wilcoxon two sample test p= 1.45^−11^) and *A*24:02* (mean = 1.37, p= 1.27^−9^) associated SNPs have significantly higher iHS values than the chromosome average, with *A*11:01* having a mean iHS that is above the 95^th^ percentile. Haplotypes carrying *HLA-A*11:01, A*24:02, B*46:01* or *B*58:01* were previously identified to have unusually high LD in this population (Chen et al. 2016). This analysis identified two other *HLA class I* alleles as having highly distinct signatures of directional selection, A*30 and C*07. Whereas HLA-C*07 is known to interact strongly with KIR to educate NK cells (Hilton et al. 2015a; Yawata et al. 2006), HLA-A*30 does not possess a KIR ligand. Together, these findings thus illustrate that HLA class I in the Chinese Southern Han has been targeted by natural selection and suggest that one major benefit has been to increase the number of KIR ligands present in the population.

We next examined whether the observed distributions of *HLA class I* encoded KIR ligands were consistent with modern human population dispersal. Cluster analysis shows there are five groups of *HLA class I* frequency spectra that correspond to the broad population groups of African, European, Asian, Oceanian and American origin (Figure 3D). By contrast, three distinct and strongly supported groups cluster according to their proportions of haplotypes encoding one, two or three KIR ligands (Figure 3E). Notable examples are the Ryukyu Japanese and Indigenous Australian populations, who group with Asian populations when analyzed by *HLA class I* haplotype distribution (Figure 3D). By contrast, Ryukyu Japanese and Australians appear more similar to Africans and other groups when analyzed according to the number of KIR ligands encoded by their *HLA class I* haplotypes (Figure 3E). In a counter example, Northern Indian and Tuvan populations (Figure S4C) cluster with Europeans when analyzed by their *HLA class I* alleles, but with East Asians when analyzed by the number of KIR ligands. Thus, *HLA class I* allele frequency distributions are consistent with the origins of the populations studied and with the pattern of human dispersal out of Africa (Henn et al. 2012), whereas the number of KIR ligands encoded by *HLA class I* haplotypes is not. This finding suggests that the similar number of KIR ligands observed across populations is likely due to convergent evolution, because distinct *HLA class I* haplotypes produce similar distributions of KIR ligands. The findings thus also support our assessment that the unusual distribution of KIR ligands in East Asian populations is due to natural selection.

In summary, these results show that similar quantities of KIR ligands can be obtained using different subsets of *HLA class I* haplotypes, indicating there is pressure to maintain a certain balance of KIR ligands across populations, regardless of the background HLA allotype, and that this balance is perturbed in East Asia. Our observations show that successive rounds of admixture followed by natural selection favouring specific *HLA class I* haplotypes have increased the quantity of KIR/HLA interactions of populations in East Asia. To investigate the characteristics of this receptor and ligand diversity, we next studied the *KIR* locus in the Chinese Southern Han.

### High frequency of inhibitory KIR allotypes in Southern Han

The *KIR* locus comprises genes encoding the four inhibitory and six activating KIR known to bind polymorphic HLA class I ligands, and three that do not bind polymorphic HLA class I (Guethlein et al. 2015). In total, we identified 116 *KIR* alleles, representing 101 KIR allotypes (Figure S5). A total of 46 novel *KIR* alleles (39.7% of total *KIR* alleles detected) were characterized (Figure S6) and 24.8% of the individual Han carried at least one novel allele. Correcting for the number of individuals tested showed that the Southern Han are more diverse than Amerindians and Oceanians, but less diverse than Europeans and Africans (Figure 4A). *KIR* diversity of the Chinese Southern Han is thus consistent with genome-wide diversity when compared to other populations (Campbell and Tishkoff 2008). The Chinese Southern Han have 70 centromeric and 91 telomeric *KIR* haplotype motifs that combine to form a minimum of 199 *KIR* haplotypes (Figure S7A-C). The majority are *KIR A* haplotypes (74.7%, Figure 4B), including 8 of the 10 most frequent haplotypes (Figure 4C). This skewing towards *KIR A* haplotypes is more pronounced in the centromeric region (87.9%) than the telomeric (79.7%) region (Figure 4C, Figure S7A-B).

**Figure 4.**
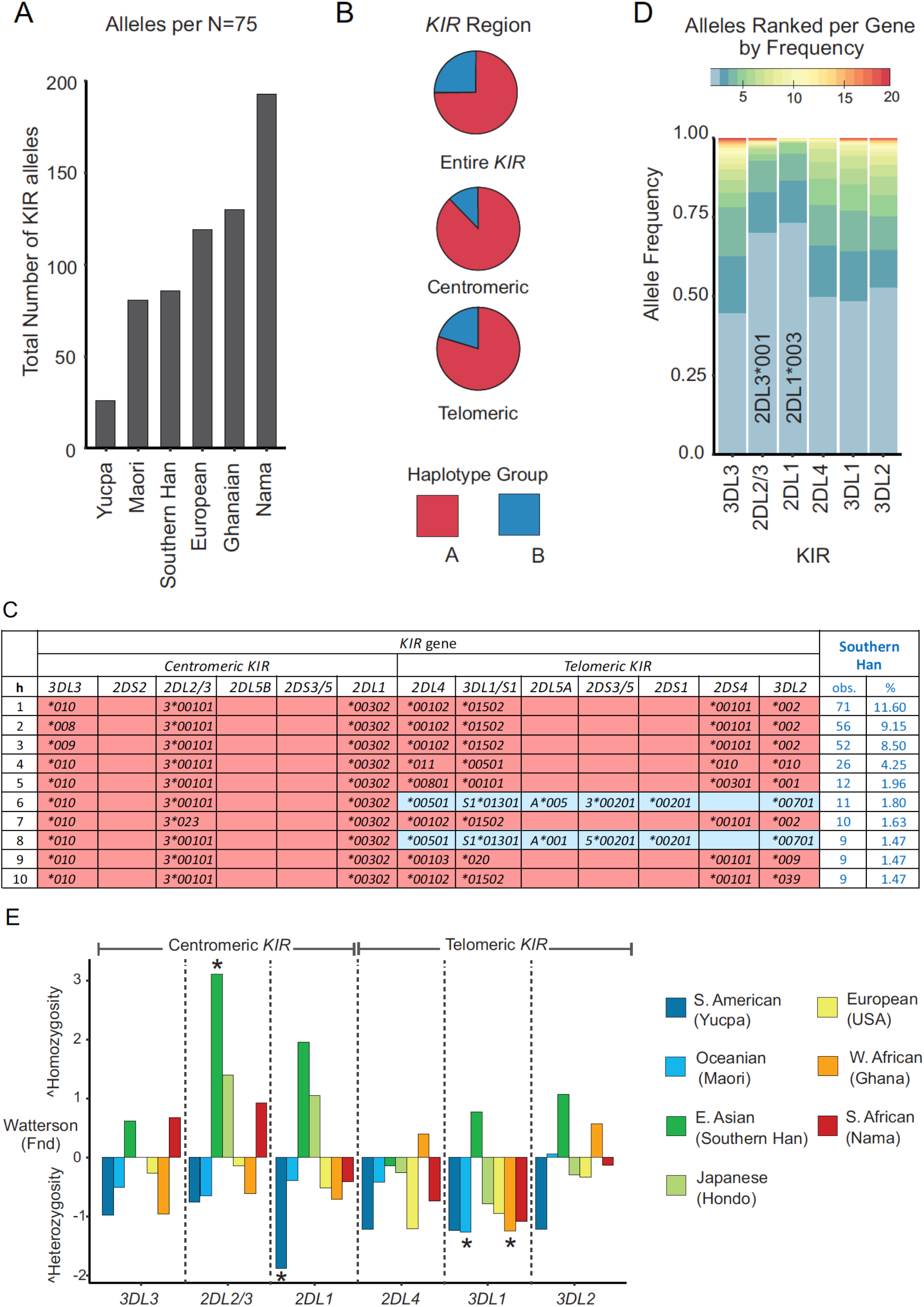
Directional selection on *centromeric KIR* in Southern Han. A. Shows the number of *KIR* alleles present in the Southern Han compared with other populations analyzed at comparable resolution; 75 individuals were selected at random from each population. B. Shown are the combined frequencies of *KIR A* (red) and *KIR B* (blue) haplotypes, for the complete haplotypes, and for the *centromeric* and *telomeric* regions. C. Shown are the ten most frequent complete, high-resolution, *KIR* haplotypes identified in the Chinese Southern Han population. *KIR A* haplotypes are shaded in red, *KIR B* haplotypes are shaded in blue. At the right is shown for each haplotype the number of individuals carrying the haplotype, and its frequency. All the haplotypes are shown in Figure S7. D. Bar graph shows a summary *KIR* allele frequencies. The colors from blue to red correspond to the rank in frequency from highest (blue) to lowest (red). Full frequency distributions are shown in Figure S5. E. Shown are normalized deviate values of Ewens-Watterson’s F test (F_nd_) in representative global populations. Positive values of Fnd indicate homozygosity, negative values indicate heterozygosity. An asterisk denotes significance (P < 0.05 or > 0.95) using the exact test (Salamon et al. 1999).

*KIR A* haplotypes encode all four inhibitory receptors that bind HLA class I ligands and either one or no activating receptors (Wilson et al. 2000). Accordingly, among the *KIR* alleles identified in the Chinese Southern Han, we observed high frequencies of those encoding strong inhibitory receptors. Both KIR2DL1*003, a strongly-inhibiting allotype of KIR2DL1 (Bari et al. 2009; Hilton et al. 2015a), and KIR2DL3*001, a strongly-inhibiting allotype of KIR2DL2/3 (Yawata et al. 2006), are common in the Chinese Southern Han, having frequencies of 73.5% and 71.1% respectively (Figure S5). Also frequent are KIR3DL1*015, which is a strong inhibitor on binding to the Bw4 ligand (Yawata et al. 2006), and KIR3DL2*002 that has high expression but unknown functional properties (Figure S5). Noticeably scarce are inhibitory KIR allotypes having mutations that prevent cell surface expression, of which there are many examples (Bari et al. 2009; Hilton et al. 2015b; Pando et al. 2003; VandenBussche et al. 2006). For instance, weakly-expressed KIR3DL1*004 is frequent in many populations (Norman et al. 2007), but absent from the Chinese Southern Han (Figure S5, and ref (Tao et al. 2014)). Also rare in the Chinese Southern Han are alleles encoding inhibitory allotypes of reduced function, such as KIR2DL1*004 (3.6%, Figure S5), which is common in other population groups (Bari et al. 2009; Meenagh et al. 2008; Nemat-Gorgani et al. 2014; Norman et al. 2013; Vierra-Green et al. 2012). Moreover, the frequencies of alleles encoding activating receptors are much lower (4.4%-18%) than those encoding inhibitory receptors (91.5%-100%), an effect compounded by presence of multiple non-functional activating KIR allotypes (Figure S5). Exceptional is KIR2DS4, for which the frequencies of functional and non-functional allotypes are balanced (55%:45%, Figure S5). These observations point to a strong requirement in the Southern Han population for retaining high numbers of functional inhibitory KIR, but not activating KIR.

### Directional selection reduced centromeric KIR region diversity in the Southern Han

In Chinese Southern Han, the *KIR3DL1/S1* and *KIR3DL2* genes encoding inhibitory NK cell receptors specific for polymorphic HLA class I ligands, have two or three high frequency alleles and multiple less frequent alleles (Figure S5). In contrast, *KIR2DL1* and *2DL2/3* also encode inhibitory receptors but are each dominated by one high frequency allele (Figure 4D). To explore this observation, we compared the observed homozygosity to the expected across populations representing major ancestry groups from Europe, Africa, Asia, South America, and Oceania, using the Ewens-Watterson test (Fnd). *KIR2DL1* and *KIR2DL2/3* are in the centromeric region of the *KIR* locus, whereas *KIR3DL1/S1* and *KIR3DL2* are telomeric *KIR* genes (Wilson et al. 2000). Overall the Southern Han show greater homozygosity compared to other populations, which is more pronounced among centromeric than telomeric *KIR* genes and statistically significant for *KIR2DL2/3* (Fnd = 3.1, P > 0.985, Figure 4E). This high-resolution analysis of KIR alleles, complements recent analysis of genome-wide SNP data that identified directional selection specifically in East Asian centromeric *KIR* (Augusto et al. 2019). The only other population exhibiting directional selection in the centromeric *KIR* region is the Hondo Japanese (Yawata et al. 2006). Thus, the profile observed for East Asian populations is distinct from other populations. Together, these analyses suggest that directional selection reduced sequence diversity of the centromeric *KIR* in the Southern Han, whereas the telomeric *KIR* region retains some diversity. In addition, we observed a minimum of eleven different *KIR* haplotypes having a duplication in the telomeric region (Figure S7D). The telomeric *KIR* have greater allelic diversity than centromeric *KIR* in Chinese Han (Figure S7D), and these duplication haplotypes have potential to further diversify the NK cell repertoire because both allotypes of each gene are expressed (Beziat et al. 2013; Norman et al. 2009). We conclude that the centromeric *KIR* region provides consistency to Chinese Southern Han NK cell receptors, whereas the telomeric *KIR* region provides NK cell receptor diversity.

### Interactions of KIR with HLA class I

NK cell function is modulated by interactions between KIR and their cognate ligands, HLA class I molecules. While all HLA-C molecules are always ligands for KIR, only a sub-set of HLA-A and -B molecules function as KIR ligands. We examined the impact of genetic variation on the diversity and quantity of KIR/HLA class I interactions in the Chinese Southern Han. Individuals have a mean of 6.7 different pairs of interacting KIR and HLA class I ligands. These form a normal distribution in which individuals have from one to twelve interactions (Shapiro -Wilk test, p = 0.147, Figure 5A). Such normal distributions are seen in other populations (Nemat-Gorgani et al. 2014; Norman et al. 2013). To investigate the distinct HLA-A and -B ligand distribution of the Southern Han we divided this analysis into its major components of KIR interactions with HLA-C, and of KIR interactions with HLA-A and -B (Figure S2). In analyzing only the interactions with HLA-C, we find that functional diversity, as measured by the mean number of different receptor/ligand combinations per individual, is consistent with the overall genetic diversity of the populations studied. At the low end of the range are the Yucpa Amerindians with two different receptor/ligand interactions per individual. At the high end are the Southern African Nama with 4.5 different interactions (Figure 5B).

**Figure 5.**
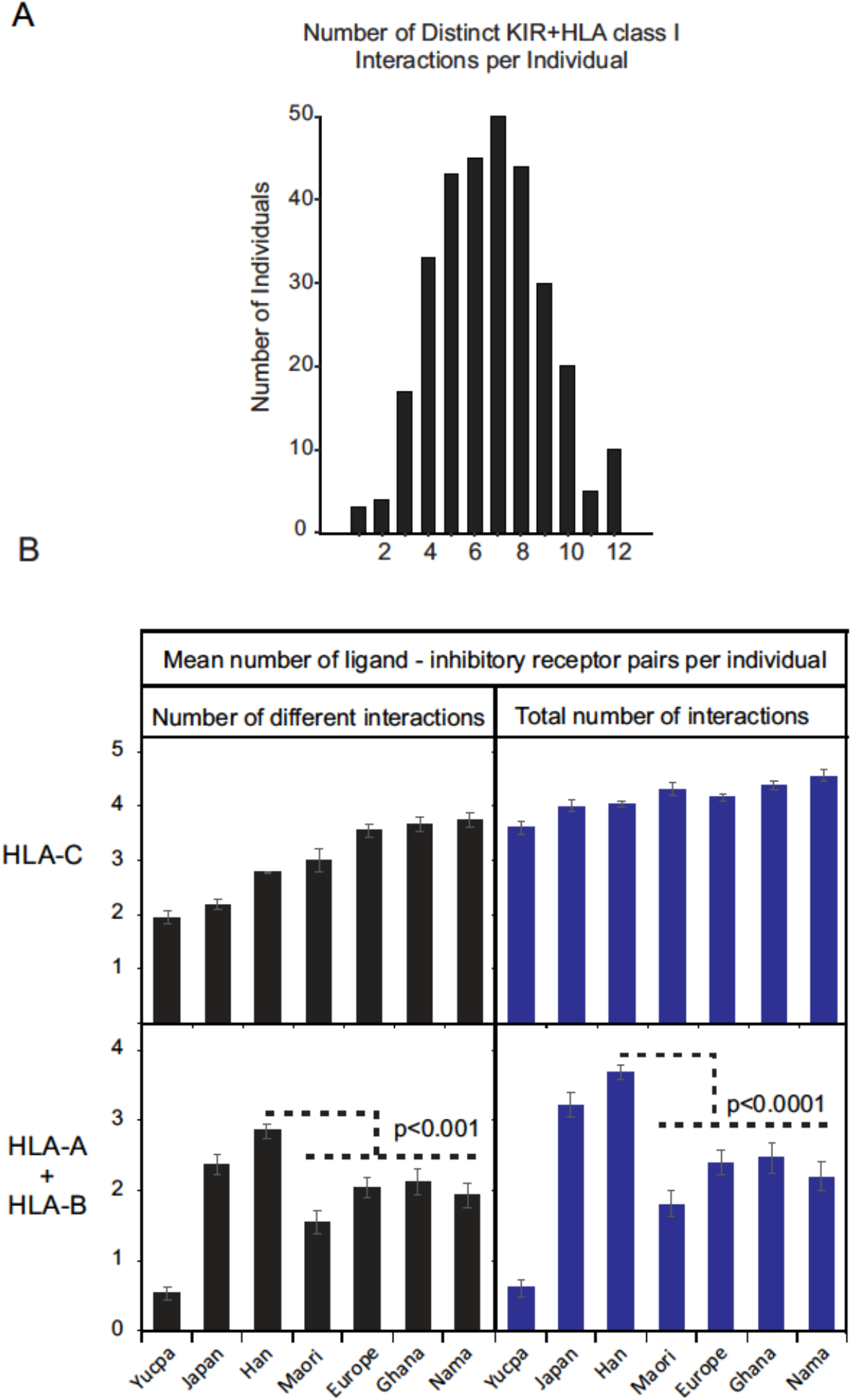
East Asians have a greater diversity of KIR interactions with HLA-A and -B than other populations. A. Plot of the number of different interacting ligand/receptor allotype pairs observed per individual in the Southern Han. B. Shows the mean number of different ligand/receptor allotype pairs per individual (left) and the mean total number of ligand/receptor allotype pairs per individual (right) for HLA-C (upper) and HLA-A and -B combined (lower). In the populations shown, *KIR* and *HLA class I* have been analyzed to similar high-resolution as described here for Southern Han. These populations comprise the Yucpa (Gendzekhadze et al. 2009), Japanese (Yawata et al. 2006), Māori (Nemat-Gorgani et al. 2014), European (Vierra-Green et al. 2012), Ghanaian (Norman et al. 2013) Nama (Nemat-Gorgani et al. 2018). Error bars are s.e.m. and p values are from a t-test.

When the total number of interacting pairs of inhibitory KIR and HLA-C ligands is analyzed, the ranking remains the same but the difference across populations is reduced, ranging from 3.6 to 4.9 viable inhibitory KIR/HLA-C interactions per individual (Figure 5B). On this scale, the Chinese Southern Han are seen to have relatively low diversity and a similar number of interactions between inhibitory KIR and HLA-C to other populations. In sharp contrast, the Chinese Southern Han, together with the Hondo Japanese, have significantly higher number (t-test, p<0.0001) and diversity (p<0.001), of inhibitory KIR interactions with HLA-A and -B than any other population (Figure 5B). Thus, both the quantity and quality of interactions between inhibitory KIR and HLA-A and -B are enhanced in Southern Han and Hondo Japanese. We predict this will also be true for other East Asian populations.

## Discussion

Our analysis shows that the geographic distribution of *HLA class I* alleles and haplotypes is consistent with human dispersal out of Africa and the distance of their geographical location from Africa. Despite the significant differences across human populations in the distributions of *HLA class I* alleles (Meyer et al. 2007), the distribution of KIR ligands is very similar. For *HLA-C*, all haplotypes encode one of two alternative KIR ligands, whereas for *HLA-A* and *HLA-B* there is a balance between haplotypes that either encode one or two KIR ligands, and haplotypes that encode no KIR ligand (Guethlein et al. 2015). Thus, a similar balance of KIR ligands is independently maintained in different human populations using very different HLA-A and -B allotypes. In sharp contrast, the Southern Han Chinese and other East Asian populations do not follow these patterns. The high frequencies of the *HLA class I* haplotypes shared by East Asian populations are an indication of their recent shared ancestry (Abdulla et al. 2009). We find a greater abundance of HLA-A and -B KIR ligands in East Asians than other populations, as well as a greater diversity of interactions between inhibitory KIR and HLA-A and -B. That our comparison included Southern African KhoeSan, whose genetic diversity is the highest among modern humans (Henn et al. 2011; Tishkoff et al. 2009), strongly suggests the high frequency and diversity of KIR ligands in East Asia is the result of natural selection.

The frequency of East Asian *HLA class I* alleles that derive from ancient humans by introgression was previously estimated to be 70-80% (Abi-Rached et al. 2011). The most common of these alleles are *HLA-A*11* and *HLA-A*24*, which encode KIR ligands. We show that more recently, *HLA-B*46:01* and *HLA-B*58:01*, which also encode KIR ligands, were specifically enhanced in frequency in Southern Han following admixture with local populations. HLA-B*46 is a good educator of NK cells (Yawata et al. 2006) and rose in frequency in South East Asia under positive selection (Abi-Rached et al. 2010). The haplotype that encodes HLA-B*58 likely arose in Northern Asia and although the signal is weaker, this may have been selected both in the Northern and Southern Han. Consequently, *HLA-B*46:01* and *HLA-B*58:01* are the most frequent *HLA-B* alleles encoding KIR ligands, and distinguish the most frequent *HLA class I* haplotypes, in the Southern Han. Together with clear demonstration of natural selection recently targeting the *MHC* region, these findings all support the proposition that natural selection in East Asia favors *HLA class I* haplotypes carrying more than one KIR ligand and suggests there were two major waves of adaptive introgression involving these haplotypes. There is evidence for adaptive introgression of *HLA* alleles in other modern human populations (Busby et al. 2017; Rishishwar et al. 2015). For example, Bantu speakers from western central Africa expanded through new habitats and acquired *HLA* haplotypes from rainforest hunter-gatherer pygmies (Patin et al. 2017). Our findings may fit with recent work identifying a second wave of Denisovan-like admixture that is specific to East Asian populations (Browning et al. 2018). Thus, although we show the *HLA-B*46:01* and -*B*58:01* haplotypes were obtained by the Han from neighboring modern human populations, they were likely to have been acquired by those populations as a consequence of admixture with archaic humans.

Complementing the high number of HLA class I ligands, we find that in the Chinese Southern Han the number of inhibitory KIR is increased relative to other groups. These KIR allotypes are distinguished by their high expression, inhibitory strength and fine specificity for ligand (Boudreau et al. 2017; Hilton et al. 2015a; Saunders et al. 2016). Possessing higher numbers of inhibitory KIR leads to better effector function, and a higher number of inhibitory KIR ligands leads to larger numbers of circulating NK cells, stronger killing and greater diversity of the NK cell repertoire (Beziat et al. 2013; Brodin et al. 2009; Yawata et al. 2006). That the number of receptors (Pelak et al. 2011) and ligands (Thons et al. 2017) correlates with infection control, suggests the diverse NK cell repertoires of the Southern Han have likely evolved to combat infectious diseases common or endemic to East Asia. Although it is difficult to identify the specific pathogen exposure history of the Chinese Southern Han, the most plausible candidates for causing selection pressure are viral infections that have established roles for KIR/HLA interaction during host defense (Abi-Rached et al. 2010; Bashirova et al. 2006). Such pathogens have been shown to be effective drivers of adaptive introgression and natural selection in human populations (Enard and Petrov 2018; Harrison et al. 2019). One example is nasopharyngeal carcinoma (NPC) caused by Epstein-Barr virus. HLA-A*11 offers protection from NPC (Tang et al. 2012), and the interaction of KIR3DL2 with HLA-A*11 is dependent on presentation of peptides derived from EBV (Hansasuta et al. 2004). Influenza is another key candidate, with highly virulent epidemics linked to the combination of dense population, agriculture and industrialization (Cao et al. 2009; Chen et al. 2006). Human specific viral hepatitis infections and arboviruses are also endemic to China and South East Asia, including Japanese encephalitis, Dengue and chikungunya (Bashirova et al. 2006; Khakoo et al. 2004; Naiyer et al. 2017; Petitdemange et al. 2011; Thons et al. 2017; Townsley et al. 2016). Consistent with these observations, *KIR A* has established roles in controlling virus infections (Bashirova et al. 2006; Khakoo et al. 2004), and we recently showed *KIR A* homozygosity protects from leukemia (Deng et al. 2019). Reproduction is also a major driver of selection, where *KIR AA*/*C2*^+^*HLA-C* genotype is associated with preeclampisia (Parham and Moffett 2013). Thus, the low frequency of C2^+^HLA in East Asia (Figure S3) likely allows the *KIR A* haplotype to reach high frequency (Nemat-Gorgani et al. 2018). High resolution analysis of KIR and HLA diversity will be critical for understanding these and other complex diseases.

In conclusion, our high-resolution analysis of KIR and HLA class I combinatorial diversity has uncovered a distinctive enhancement of the interactions between inhibitory KIR and HLA-A and -B in East Asians. These genetically determined distinctions likely underlie differences across human populations in their susceptibility to infections and immune-mediated diseases.

## Supporting information

Figure S2 KIR and HLA Interactions

Figure S4 HLA Haplotype Frequencies

Figure S7 KIR Haplotype Frequencies

Supplemental Figures With Text

## Supplemental Data

Supplemental Data include four figures and three Excel spreadsheets.

## Acknowledgements

This study was supported by the Science, Technology and Innovation Commission of Shenzhen Municipality (grant number: JCYJ20190806152001762 to ZD), the National Natural Science Foundation of China (grant number: 81373158 to ZD) and the National Institutes of Health of the USA (grant numbers: NIH R01 AI017892 to PP, and R56 AI151549 to PJN). We thank the Chinese blood donors for generously providing DNA samples for this study.

## Web Resources and Accession Numbers

The URLs for data, material and programs used herein are as follows: The scripts used in the study are located at https://github.com/n0rmski/Han_Study/

ImmunoPolymorphism database (IPD), http://www.ebi.ac.uk/ipd/

International Histocompatibility Working Group (IHWG), www.ihwg.org/

The official IPD names (Robinson et al. 2015) and GenBank accession numbers for the *KIR* sequences reported in this paper are: (*KIR* prefix excluded for brevity) *2DL1*00304* (KT438851), *2DL1*00305* (KT438852), *2DL1*030* (KP025959), *2DL1*031* (KP025960), *2DL1*033* (KT438853), *2DL1*034* (KT438854), *2DL2*013* (KM017076), *2DL3*00109* (KF766495), *2DL3*00110* (KF766497), *2DL3*025* (KF766496), *2DL3*026* (KF766498), *2DL3*027* (KF766499), *2DL3*028* (KF766500), *2DL3*029* (KF766501), *2DL3*031* (KF849247), *2DL4*00503* (KT438855), *2DL4*00504* (KT438856), *2DL4*032* (KT438858), *2DL4*033* (KT438859), *2DL4*034* (KT438857), *2DL5A*022* (KT438863), *2DS2*009* (KT438862), *2DS4*00105* (KP025962), *2DS4*017* (KP025961), *2DS4*018* (KP025963), *3DL1*01505* (KF849249), *3DL1*079* (KF849250), *3DL2*00706* (KT899864), *3DL2*00707* (KT899868), *3DL2*083* (KT899867), *3DL2*084* (KT438861), *3DL2*091* (KT438860), *3DL2*093* (KT899866), *3DL2*099* (KT899865), *3DL3*01003* (KU529275), *3DL3*02602* (KU529271), *3DL3*04802* (KU529269), *3DL3*062* (KU529272), *3DL3*063* (KU529270), *3DL3*064* (KU529273), *3DL3*065* (KU529274), *3DS1*078* (KJ001806), *3DS1*082* (KJ001804), *3DS1*083* (KJ001805), *3DS1*084* (KJ001807), *3DS1*085* (KJ365317).

## Notes

### Competing Interest Statement

The authors have declared no competing interest.

### Summary of Updates

A typo on the X-axis of Figure 2A was corrected.

