## Supplemental Figures With Text for "Genetically Determined Strength of Natural Killer Cells is Enhanced by Adaptive HLA class I Admixture in East Asians"

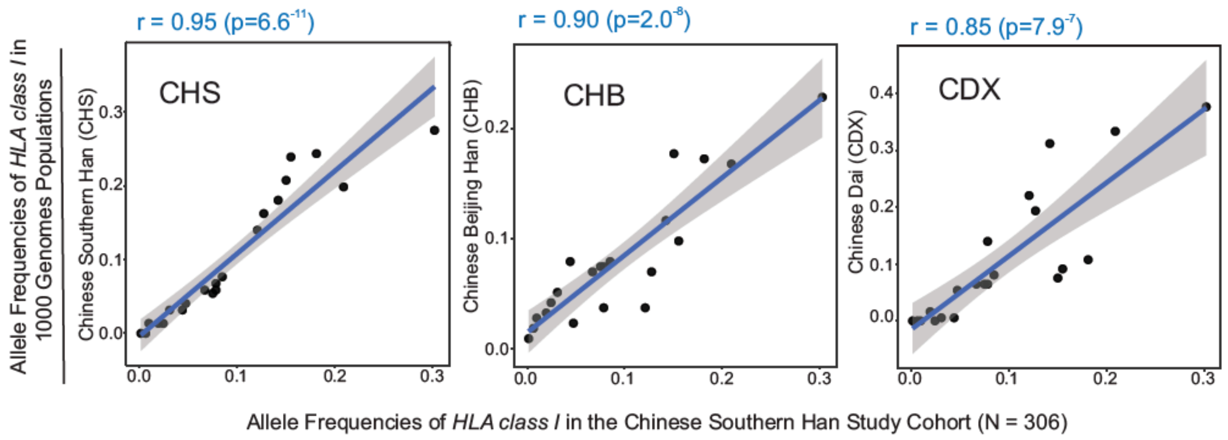
r = 0.95 (p=6.6-11)
r = 0.90 (p=2.0-8)
r = 0.85 (p=7.9-7)
Allele Frequencies of HLA class I in 1000 Genomes Populations
Chinese Southern Han (CHS)
Chinese Beijing Han (CHB)
Chinese Dai (CDX)
CHS
CHB
CDX
Allele Frequencies of HLA class I in the Chinese Southern Han Study Cohort (N = 306)

**Figure S1**

| HLA-A | | HLA-B | | | HLA-C | |
|---|---|---|---|---|---|---|
| Allele | Frequency (%) | Allele | -21M/T | Frequency (%) | Allele | Frequency (%) |
| *A*01:01* | 2.45 | *B*07:02* | M | 0.98 | *C*01:02* | 20.92 |
| *A*01:03* | 0.16 | *B*07:05* | M | 0.49 | *C*01:03* | 0.33 |
| *A*02:01* | 8.82 | *B*08:01* | M | 0.65 | *C*02:02* | 0.16 |
| *A*02:03* | 5.39 | *B*13:01* | T | 7.84 | *C*03:01* | 0.16 |
| *A*02:05* | 0.33 | *B*13:02* | T | 2.12 | *C*03:02* | 7.52 |
| *A*02:06* | 2.45 | *B*14:02* | M | 0.16 | *C*03:03* | 4.58 |
| *A*02:07* | 12.26 | *B*15:01* | T | 3.6 | *C*03:04* | 12.75 |
| *A*02:17* | 0.16 | *B*15:02* | T | 4.74 | *C*03:17* | 0.16 |
| *A*03:01* | 1.14 | *B*15:05* | T | 0.49 | *C*03:38* | 0.16 |
| *A*03:02* | 0.16 | *B*15:07* | T | 0.16 | *C*04:01* | 4.58 |
| *A*11:01* | 30.23 | *B*15:11* | T | 1.14 | *C*04:03* | 1.8 |
| *A*11:02* | 2.78 | *B*15:12* | T | 0.49 | *C*05:01* | 0.49 |
| *A*11:03* | 0.16 | *B*15:18* | T | 0.65 | *C*06:02* | 4.41 |
| *A*11:188* | 0.16 | *B*15:21* | T | 0.16 | *C*07:01* | 0.49 |
| *A*23:01* | 0.33 | *B*15:25* | T | 0.49 | *C*07:02* | 18.14 |
| *A*24:02* | 15.19 | *B*15:27* | T | 0.33 | *C*07:04* | 0.65 |
| *A*24:10* | 0.16 | *B*18:01* | T | 0.33 | *C*07:06* | 0.33 |
| *A*26:01* | 1.96 | *B*27:04* | T | 1.31 | *C*07:19* | 0.49 |
| *A*29:01* | 0.65 | *B*27:05* | T | 0.16 | *C*07:43* | 0.16 |
| *A*30:01* | 2.45 | *B*27:06* | T | 0.16 | *C*08:01* | 8.5 |
| *A*31:01* | 3.11 | *B*35:01* | T | 2.29 | *C*08:02* | 0.16 |
| *A*32:01* | 1.47 | *B*35:02* | T | 0.16 | *C*08:03* | 0.49 |
| *A*33:01* | 0.33 | *B*35:03* | T | 0.33 | *C*12:02* | 2.94 |
| *A*33:03* | 6.86 | *B*37:01* | T | 1.14 | *C*12:03* | 2.12 |
| *A*34:01* | 0.16 | *B*38:01* | M | 0.16 | *C*14:02* | 3.43 |
| *A*68:01* | 0.49 | *B*38:02* | M | 2.94 | *C*14:03* | 0.16 |
| *A*74:02* | 0.16 | *B*39:01* | M | 1.96 | *C*15:02* | 3.11 |
| | | *B*39:05* | M | 0.16 | *C*15:05* | 0.49 |
| | | *B*40:01* | T | 15.52 | *C*16:02* | 0.16 |
| | | *B*40:02* | T | 1.31 | *C*17:01* | 0.16 |
| | | *B*40:06* | T | 1.96 | | |
| | | *B*40:40* | T | 0.16 | | |
| | | *B*41:01* | T | 0.16 | | |
| | | *B*44:02* | T | 0.65 | | |
| | | *B*44:03* | T | 0.65 | | |
| | | *B*44:07* | T | 0.16 | | |
| | | *B*46:01* | T | 14.22 | | |
| | | *B*48:01* | M | 0.98 | | |
| | | *B*48:03* | M | 0.82 | | |
| | | *B*49:01* | T | 0.16 | | |
| | | *B*50:01* | T | 0.16 | | |
| | | *B*51:01* | T | 3.76 | | |
| | | *B*51:02* | T | 1.31 | | |
| | | *B*52:01* | T | 2.12 | | |
| | | *B*54:01* | T | 4.09 | | |
| | | *B*55:01* | T | 0.16 | | |
| | | *B*55:02* | T | 4.09 | | |
| | | *B*56:01* | T | 1.63 | | |
| | | *B*56:04* | T | 0.16 | | |
| | | *B*57:01* | T | 1.14 | | |
| | | *B*58:01* | T | 7.84 | | |
| | | *B*59:01* | T | 0.16 | | |
| | | *B*67:01* | M | 0.65 | | |
| | | *B*81:02* | M | 0.33 | | |

KIR ligand

- A3/11
- Bw4
- C1
- C2

Fig

**Figure S3**

### Centromeric *KIR*

| 3DL3 | Freq. |
|---|---|
| *010 | 0.448 |
| *008 | 0.180 |
| *009 | 0.155 |
| *006 | 0.046 |
| *001 | 0.041 |
| *015 | 0.033 |
| † *04802 | 0.020 |
| *028 | 0.018 |
| † *02602 | 0.011 |
| *048 | 0.011 |
| *004 | 0.008 |
| *002 | 0.005 |
| *003 | 0.005 |
| † *062 | 0.005 |
| † *064 | 0.005 |
| † *063 | 0.003 |
| † *065 | 0.003 |
| † *01003 | 0.002 |
| *013 | 0.002 |

| 2DS2 | Freq. |
|---|---|
| *00101 | 0.111 |
| † *009 | 0.005 |
| neg | 0.884 |

| 2DL5B | Freq. |
|---|---|
| B*006 | 0.042 |
| B*010 | 0.023 |
| B*002 | 0.016 |
| B*008 | 0.002 |
| neg | 0.917 |

| 2DS3/5 | Freq. |
|---|---|
| 5*00201 | 0.044 |
| 3*001 | 0.033 |
| 3*00201 | 0.008 |
| neg | 0.915 |

| 2DL2/3 | Freq. |
|---|---|
| 3*00101 | 0.701 |
| 3*00201 | 0.129 |
| 2*00301 | 0.100 |
| 3*023 | 0.021 |
| 2*00101 | 0.018 |
| † 3*00109 | 0.007 |
| 3*015 | 0.003 |
| 3*022 | 0.003 |
| † 3*028 | 0.003 |
| † 3*00110 | 0.002 |
| 3*019 | 0.002 |
| 3*021 | 0.002 |
| † 3*025 | 0.002 |
| † 3*026 | 0.002 |
| † 3*027 | 0.002 |
| † 3*029 | 0.002 |
| † 3*031 | 0.002 |
| † 2*013 | 0.002 |

| 2DL1 | Freq. |
|---|---|
| *00302 | 0.735 |
| *00201 | 0.134 |
| *00401 | 0.036 |
| *001 | 0.002 |
| † *00304 | 0.002 |
| † *00305 | 0.002 |
| † *030 | 0.002 |
| † *031 | 0.002 |
| † *033 | 0.002 |
| † *034 | 0.002 |
| neg | 0.085 |

### Telomeric *KIR*

| 2DL4 | Freq. |
|---|---|
| *00102 | 0.498 |
| *00501 | 0.163 |
| *011 | 0.129 |
| *006 | 0.082 |
| *00801 | 0.052 |
| *00103 | 0.044 |
| † *00503 | 0.005 |
| † *00504 | 0.003 |
| † *032 | 0.002 |
| † *033 | 0.002 |
| † *034 | 0.002 |
| neg | 0.018 |

| 3DL1 | Freq. |
|---|---|
| *01502 | 0.485 |
| S1*01301 | 0.158 |
| *00501 | 0.129 |
| *00701 | 0.083 |
| *00101 | 0.049 |
| *020 | 0.041 |
| *02901 | 0.016 |
| † *070 | 0.003 |
| † S1*082 | 0.003 |
| † S1*085 | 0.003 |
| *00502 | 0.002 |
| *008 | 0.002 |
| † *01505 | 0.002 |
| † *079 | 0.002 |
| † S1*078 | 0.002 |
| † S1*083 | 0.002 |
| † S1*084 | 0.002 |
| neg | 0.016 |

| 2DL5A | Freq. |
|---|---|
| A*001 | 0.114 |
| A*005 | 0.052 |
| A*012 | 0.007 |
| † A*022 | 0.002 |
| neg | 0.825 |

| 2DS3/5 | Freq. |
|---|---|
| 5*00201 | 0.123 |
| 3*00201 | 0.049 |
| neg | 0.828 |

| 2DS4 | Freq. |
|---|---|
| *00101 | 0.542 |
| *010 | 0.127 |
| *00401 | 0.083 |
| *00301 | 0.052 |
| † *018 | 0.003 |
| † *00105 | 0.002 |
| † *017 | 0.002 |
| neg | 0.188 |

| 2DS1 | Freq. |
|---|---|
| *00201 | 0.175 |
| *00202 | 0.003 |
| *006 | 0.002 |
| neg | 0.820 |

| 3DL2 | Freq. |
|---|---|
| *002 | 0.528 |
| *010 | 0.119 |
| *00701 | 0.108 |
| *008 | 0.067 |
| *001 | 0.057 |
| *009 | 0.036 |
| *039 | 0.031 |
| *015 | 0.010 |
| *021 | 0.010 |
| † *093 | 0.010 |
| *016 | 0.005 |
| † *091 | 0.005 |
| † *00706 | 0.003 |
| † *00707 | 0.003 |
| *027 | 0.003 |
| † *083 | 0.002 |
| † *084 | 0.002 |
| † *099 | 0.002 |

**Figure S5**

| Novel allele name | GenBank Accession | Most similar allele | Nucleotide change (position in full CDS) | Exon | Substitution (Codon Number) | Amino acid substitution | ref |
|---|---|---|---|---|---|---|---|
| *2DL1*00304* | KT438851 | *2DL1*00302* | 582 G > A | 5 | 173 GGG > GGA | syn | |
| *2DL1*00305* | KT438852 | *2DL1*00302* | 1044 A > G | 9 | 327 CCA > CCG | syn | |
| *2DL1*030* | KP025959 | *2DL1*00302* | 343 A > G | 4 | 94 AGT > GGT | Ser > Gly | a |
| *2DL1*031* | KP025960 | *2DL1*00302* | 188 A > G | 4 | 42 GAG > GGG | Glu > Gly | b |
| *2DL1*033* | KT438853 | *2DL1*00302* | 867 C > G | 8 | 268 AGC > AGG | Ser > Arg | c |
| *2DL1*034* | KT438854 | *2DL1*00302* | 13 G > T | 1 | 17 GTC > TTC | Val > Phe | d |
| *2DL2*013* | KM017076 | *2DL2*00302* | 806 C > G | 7 | 248 TCC > TGC | Ser > Cys | e |
| | | | 1018 T > C | 9 | 319 TCC > CCC | Ser > Pro | |
| *2DL3*00109* | KF766495 | *2DL3*00101* | 478 C > T | 5 | 139 CTA > TTA | syn | e |
| *2DL3*00110* | KF766497 | *2DL3*00101* | 618 A > C | 5 | 185 CCA > CCC | syn | f |
| *2DL3*025* | KF766496 | *2DL3*00101* | 280 C > A | 4 | 73 CTT > ATT | Leu > Ile | e |
| *2DL3*026* | KF766498 | *2DL3*00101* | 202 G > A | 4 | 47 GAC > AAC | Asp > Asn | e |
| *2DL3*027* | KF766499 | *2DL3*00101* | 809 G > C | 7 | 249 TGC > TCC | Cys > Ser | e |
| *2DL3*028* | KF766500 | *2DL3*00101* | 800 G > A | 7 | 246 CGC > CAC | Arg > His | e |
| *2DL3*029* | KF766501 | *2DL3*00101* | 505 C > A | 5 | 148 CGT > AGT | Arg > Ser | e |
| *2DL3*031* | KF849247 | *2DL3*00101* | 735 T > C | 7 | 224 CAT > CAC | syn | e |
| | | | 736 G > A | 7 | 225 GTT > ATT | Val > Ile | |
| *2DL4*00503* | KT438855 | *2DL4*00501* | 888 G > A | 9 | 273 CAG > CAA | syn | g |
| *2DL4*00504* | KT438856 | *2DL4*00501* | 987 G > T | 9 | 306 GTG > GTT | syn | g |
| *2DL4*032* | KT438858 | *2DL4*00102* | 223 A > G | 3 | 52 AAC > GAC | Asn > Asp | g |
| *2DL4*033* | KT438859 | *2DL4*00102* | 1012 C > T | 9 | 315 CCC > TCC | Pro > Ser | g |
| *2DL4*034* | KT438857 | *2DL4*00501* | 200 G > T | 3 | 44 GGG > GTG | Gly > Val | g |
| *2DL5A*022* | KT438863 | *2DL5A*00501* | 289 G > A | 3 | 76 GGT > AGT | Gly > Ser | |
| *2DS2*009* | KT438862 | *2DS2*00101* | 109 C > T | 4 | 16 CCC > TCC | Pro > Ser | |
| *2DS4*00105* | KP025962 | *2DS4*00101* | 657 C > T | 5 | 198 TCC > TCT | syn | h |
| *2DS4*017* | KP025961 | *2DS4*00101* | 707 C > G | 6 | 215 TCC > TGC | Ser > Cys | h |
| *2DS4*018* | KP025963 | *2DS4*010* | 316 C > G | 4 | 85 CAC > GAC | His > Asp | h |
| *3DL1*01505* | KF849249 | *3DL1*01502* | 906 C > T | 5 | 281 TAC > TAT | syn | i |
| *3DL1*079* | KF849250 | *3DL1*01502* | 1119 G > T | 8 | 352 ATG > ATT | Met > Ile | i |
| *3DL2*00706* | KT899864 | *3DL2*00701* | 495 A > G | 4 | 144 TCA > TCG | syn | j |
| *3DL2*00707* | KT899868 | *3DL2*00701* | 783 C > A | 5 | 240 GCC > GCA | syn | j |
| *3DL2*083* | KT899867 | *3DL2*008* | 292 T > A | 3 | 77 TCA > ACA | Ser > Thr | |
| *3DL2*084* | KT438861 | *3DL2*00902* | 502 G > A | 4 | 147 GTT > ATT | Val > Ile | j |
| *3DL2*091* | KT438860 | *3DL2*01001* | 1315 A > C | 9 | 418 AAA > CAA | Lys > Gln | j |
| *3DL2*093* | KT899866 | *3DL2*00701* | 532 G > A | 4 | 157 GCC > ACC | Ala > Thr | |
| *3DL2*099* | KT899865 | *3DL2*00902* | 296 G > A | 3 | 78 CGC > CAC | Arg > His | |
| | | | 308 T > C | 3 | 82 CTC > CCC | Leu > Pro | |
| *3DL3*01003* | KU529275 | *3DL3*01001* | 408 G > A | 4 | 115 TCG > TCA | syn | |
| *3DL3*02602* | KU529271 | *3DL3*026* | 1074 A > G | 8 | 337 CAA > CAG | syn | |
| *3DL3*04802* | KU529269 | *3DL3*048* | 1074 A > G | 8 | 337 CAA > CAG | syn | k |
| *3DL3*062* | KU529272 | *3DL3*026* | 1074 A > G | 8 | 337 CAA > CAG | syn | |
| | | | 1184 C > T | 9 | 374 ACT > ATT | Thr > Ile | |
| *3DL3*063* | KU529270 | *3DL3*048* | 1074 A > G | 8 | 337 CAA > CAG | syn | |
| | | | 1184 C > T | 9 | 374 ACT > ATT | Thr > Ile | |
| *3DL3*064* | KU529273 | *3DL3*00601* | 1184 C > T | 9 | 374 ACT > ATT | Thr > Ile | |
| *3DL3*065* | KU529274 | *3DL3*00102* | 1184 C > T | 9 | 374 ACT > ATT | Thr > Ile | |
| *3DS1*078* | KJ001806 | *3DS1*01301* | 775 G > C | 5 | 238 GGG > CGG | Gly > Arg | i |
| *3DS1*082* | KJ001804 | *3DS1*01301* | 1114 T > C | 8 | 351 GCT > GCC | syn | i |
| *3DS1*083* | KJ001805 | *3DS1*01301* | 393 T > G | 4 | 110 GGT > GGG | syn | i |
| | | | 400 G > C | 4 | 113 GTG > CTG | Val > Leu | |
| | | | 416 G > C | 4 | 118 AGA > ACA | Arg > Thr | |
| *3DS1*084* | KJ001807 | *3DS1*01301* | 308 C > T | 3 | 82 CCC > CTC | Pro > Leu | i |
| *3DS1*085* | KJ365317 | *3DS1*01301* | 934 C > T | 5 | 291 CTT > TTT | Leu > Phe | i |

**Figure S6**

**Supplemental Material**

**Figure S1. Strong correlation of HLA alleles with CHS from 1000 Genomes data**

Shown are Pearson's correlations of frequencies for the alleles comprising the ten most frequent *HLA class I* haplotypes we observed in the Chinese Southern Han population, with those determined from the three Chinese populations available from 1000 Genomes data (Gourraud et al. 2014), CHS (Chinese Southern Han), CHB (Chinese Han Beijing) and CDX (Chinese Dai).

**Figure S2. Interactions between KIR and HLA class I allotypes**

(Excel spreadsheet)

Shown at the left are the HLA class I allotypes present in the populations analyzed for this study. Allotypes that carry one of the four alternative KIR ligands are indicated -A3/11, Bw4, C1 or C2. At the top is shown the KIR molecules that interact with these ligands. In the main body of the figure "Y" in orange shading indicates a viable KIR/HLA class I allotype interaction. For each KIR molecule the first officially named allotype (e.g. KIR2DL1*001) is indicated, and those allotypes with binding specificity differing from the first named allotype are indicated subsequently in blue text (e.g. KIR2DL1*022). In general, KIR2DL1 binds to $C2^{+}$ HLA-C allotypes, KIR2DL2 binds to $C1^{+}$ and $C2^{+}$ HLA, KIR2DL3 binds to only $C1^{+}$ HLA (Hilton et al. 2015a; Moesta et al. 2008), KIR2DS1 and some KIR2DS5 allotypes bind to $C2^{+}$ HLA-C (Blokhuis et al. 2017; Moesta et al. 2010), KIR2DS2 binds to HLA-C*16 (Liu et al. 2014). KIR2DS4 binds to HLA-A*11 and some HLA-C (Graef et al. 2009). KIR3DL1 binds to $Bw4^{+}$ HLA-A and -B (Saunders et al. 2016). KIR3DL2 binds to HLA-A*03 and -A*11 allotypes (Dohring et al. 1996; Hansasuta et al. 2004). KIR3DS1, an activating KIR3DL1/S1 allotype, was not included in the analysis because it is specific for HLA-F (Burian et al. 2016; Garcia-Beltran et al. 2016). Neither were KIR2DL4-5, KIR2DS3 or KIR3DL3 included, because they do not bind HLA-A, -B or -C (Moesta et al. 2010; VandenBussche et al. 2009)

**Figure S3. Frequencies of *HLA class I* alleles in the Chinese Southern Han population**

*HLA-A*, *-B* and *-C* allele frequencies in a panel of 306 unrelated Southern Han. Allotypes carrying one of the four alternative KIR ligands are indicated; A3/11(yellow), Bw4 (green), C1 (red) or C2 (blue). Also indicated is the M/T dimorphism at position -21 in the leader peptide of HLA-B, which favors development of NK cells that are educated by CD94:NKG2A instead of KIR (Horowitz et al. 2016) (These are rare in Chinese Southern Han, the most frequent being HLA-B*38 (3.1%), which is also a KIR ligand).

**Figure S4. Frequencies of *HLA class I* haplotypes in the Chinese Southern Han population**

(Excel spreadsheet)

A. *HLA-A-B-C* haplotype frequencies in the 306 unrelated Chinese Southern Han (2N=612). Shading indicates *HLA* alleles encoding KIR ligands. Yellow - A3/11, green - Bw4, red - C1, and blue - C2. No shading indicates the HLA allotype does not bind KIR.

B. *HLA-A-B-C* haplotypes detected in the eight representative populations shown in Figure 1. The population names are given in the top row and the number of individuals in the second row. The haplotypes and their frequencies are listed underneath each population.

C. *HLA-A-B-C* haplotypes detected in the populations shown in Figure 3.

**Figure S5. *KIR* allele frequencies**

Shown are the *KIR* allele frequencies observed in 306 unrelated Southern Han individuals (2N=612). The *KIR* genes are shown according to the orientation of the *KIR* locus segment where they are located: *Centromeric KIR* (upper) and *Telomeric KIR* (lower). Green text indicates inhibitory KIR, purple text –activating KIR. Red text indicates allotypes not expressed at the cell surface (2DS5*002 is expressed but does not bind HLA-C (Blokhuis et al. 2017)). *Neg* – gene absent. Dagger – novel allele characterized during this study.

**Figure S6. *KIR* alleles newly-identified in Chinese Southern Han**

Shown are 46 novel *KIR* alleles identified during the course of this study. From left to right: the *KIR* gene, new allele name, GenBank ID, the closest match to previously identified alleles, nucleotide changes compared to the closest match, corresponding amino acid substitutions, and frequency in the panel of 306 individuals. At the right "ref" indicates those alleles described in the following reports: a (He et al. 2017), b (Sun et al. 2016), c (Zhang et al. 2017), d (Chen et al. 2017), e (Zhen et al. 2015), f (Zhang and Deng 2017), g (Zhen et al. 2017), h (Deng et al. 2015), i (Norman et al. 2016). j (Deng et al. 2017).

**Figure S7. Frequencies of *KIR* haplotypes in the Chinese Southern Han population**

(Excel spreadsheet)

Frequencies are shown for the 306 Chinese Southern Han

A. *Centromeric KIR* haplotypes. *KIR A* haplotypes are shaded in red, *KIR B* haplotypes are shaded in blue. Yellow shading indicates novel *KIR* alleles identified in this study, which are described in Figure S4G.

B. *Telomeric KIR* haplotypes. *KIR A* haplotypes are shaded in red, *KIR B* haplotypes are shaded in blue. Yellow shading indicates novel alleles identified in this study, which are described in Figure S4G.

C. Complete *KIR* haplotypes.

D. Shows the *KIR* telomeric region haplotypes identified in the Chinese Southern Han that are characterized with duplicated *KIR2DL4* and *KIR3DL1/S1*. Eleven different haplotypes were identified, distinguished by their alleles of *KIR3DL1/S1* and other *KIR* genes, and the number of examples observed is shown at the right.
